# Stage-aware transcriptomics reveals selective haplotype persistence in short-term *ex vivo* cultured *Plasmodium vivax*

**DOI:** 10.64898/2026.05.11.724466

**Authors:** Beka Abagero, Franck Dumetz, Colby T Ford, Tirusew Tolosa, Daniel Tesefay, Biniyam Lukas, Tassew Shenkutie, Jean Popovici, Delenesaw Yewhalaw, David Serre, Eugenia Lo

**Author notes:** **Correspondence:** Eugenia Lo.

## Abstract

*Plasmodium vivax* (*Pv*) infections are developmentally asynchronous and often polyclonal, complicating interpretation of bulk parasite transcriptomes. Here, we analyzed paired *in vivo* and short-term *ex vivo* transcriptomes from Ethiopian clinical isolates using stage deconvolution and *PvMSP*1 haplotyping. *Ex vivo* maturation modestly increased inferred schizont representation while largely preserving the proportion of trophozoites and gametocytes. After adjustment for parasite stage composition, *in vivo* and *ex vivo* transcriptomes remained globally similar, with no genes significantly differentially expressed, indicating the absence of major culture-induced transcriptional response. In contrast, short-term culture reduced multiplicity of infection, contracted within-host haplotype diversity, and non-randomly depleted specific haplotypes, consistent with a clonal bottleneck. In a subset of low-complexity infections, residual expression patterns were clustered by dominant haplotype, suggesting genotype-associated transcriptional heterogeneity independent of developmental stage. Together, these findings indicate that short-term *ex vivo* culture enriches late asexual stages and selectively filters clones rather than inducing a common transcriptional program. These results shows that ex vivo cultures are reliable way to study gene expression, especially for late stages. However, these needs explicitly model developmental composition and infection complexity when interpreting *Pv* transcriptomes from natural infections

**Author summary:** Malaria caused by *Plasmodium vivax* is difficult to study because this parasite cannot yet be grown continuously in the laboratory and infections in patients often contain parasites at different developmental stages and multiple parasite lineages at the same time. In this study, we wanted to understand how much of the parasite gene-expression signal reflects true biological differences, and how much is explained by parasite development or changes that occur during short-term laboratory maturation. We compared parasites collected directly from patients in Ethiopia with matched parasite matured briefly outside the body. We found that short-term culture mainly increased the proportion of later-stage parasites, but after accounting for developmental stage, the overall gene-expression patterns remained very similar. However, culture reduced the diversity of parasite lineages within infections, suggesting that some parasite lineages survive better than others under laboratory conditions. Our findings highlight that natural *Pv* infections are complex mixtures of parasite stages and lineages. Accounting for this complexity will improve how researchers interpret parasite gene-expression studies and design future studies of parasite invasion, transmission, and survival.

## Introduction

*Plasmodium vivax* (*Pv*) is the most widespread human malaria parasite, with a global presence across Asia, Latin America, and parts of Africa. Its unique biological features, including dormant liver-stage hypnozoites, early gametocyte development, a strong preference for reticulocyte invasion, and relatively low but persistent peripheral low-density parasitemia pose major challenges to malaria elimination. In 2023, malaria caused an estimated 263 million cases and 597,000 deaths worldwide, with *Pv* contributing substantially to disease burden in co-endemic regions. Despite this public health importance, *Pv* remains comparatively understudied relative to *P. falciparum (Pf)* [1, 2], owing in part to the absence of a long-term *in vitro* culture system, low parasitemia, and asynchronous parasite stages in clinical infections that complicate the study of *Pv* gene expression and regulation [3–7].

In moderate and high transmission settings, *Pv* infections are typically asynchronous and polyclonal, comprising parasites at multiple intraerythrocytic developmental stages and harboring multiple genetic clones [3, 8, 9]. This complexity complicates efforts to resolve stage-specific gene expression programs, and to study specific biological processes (e.g., invasion-related are typically expressed late during schizogony). Consequently, the transcriptional mechanisms underlying erythrocyte invasion remain poorly understood [10–14]. Recent advances in single-cell RNA sequencing (scRNA-seq) and computational approaches have begun to overcome some of these challenges by enabling stage-specific gene expression profiling of individual parasites within asynchronous populations [7, 15–17]. However, the application of scRNA-seq in large-scale field studies is limited by factors such as logistics, high cost, challenges in removing human WBCs, parasite cell fragility, and technical complexity.

Most prior *Pv* transcriptomic studies have relied on bulk RNA-seq of either *in vivo* whole blood or short-term *ex vivo* cultured parasites [6, 14, 16]. However, there were several limitations, including small samples that could hinder the ability to confidently infer whether certain genes are consistently up- or down-regulated among genetically diverse *Pv* isolates [18–21], a lack of schizont- and gametocyte-stage specific gene data in *in vivo* samples [22] critical for reticulocyte invasion and transmission [7, 23–25], and insufficient consideration of host transcriptomic signals. While *ex vivo* cultured samples showed higher expression levels of schizont- and gametocyte-associated genes than *in vivo* whole blood samples [6, 26, 27], genes expressed by the parasites under a controlled environment may not accurately reflect the complexities of natural infections. Genes essential for schizont function may be underrepresented or absent in *in vivo* datasets, potentially leading to a skewed interpretation of gene expression dynamics [11, 26, 28, 29]. The substantial differences in gene expression across parasite stages could further underscore the complexity of *Pv* development in the human host and *ex vivo* conditions [22, 30]. These discrepancies pose challenges to the generalizability of transcriptomic findings.

Polyclonal *Pv* infections, driven by superinfection and/or relapse from hypnozoites, are common in malaria endemic countries [31, 32]. Previous longitudinal studies of *Pv* patients revealed multiple co-circulating clones within an infection based on amplicon deep sequencing (mean MOI ∼2–3; up to ≥3–6 haplotypes) and that relapses often introduce genotypes distinct from the presenting bloodstream population [33–37]. In *P. falciparum*, clonality has been shown to alter gene expression: multiple clones within an infection can modulate *var* gene expression patterns associated with antigenic variation and disease state, and longer gametocyte carriage, consistent with competition driven shifts in investment [38–40]. Prior population genetic analyses showed nonrandom clustering of parasite genotypes within polyclonal infections, implying that host-mediated selection could bias expressed phenotypes [41]. These observations in *Pf* suggest that genotype-associated expression differences in *Pv* may influence parasite survival within the host and under *ex vivo* conditions. Variation in invasion ligand repertoires, e.g., Duffy-binding protein 1 *(PvDBP1*) duplications, and sequence diversity in erythrocyte-binding protein *PvEBP/DBP2* and reticulocyte binding proteins *(RBP2b),* may offers mechanisms for clonal differences in red blood cell tropism, reticulocyte preference, and immune visibility [42, 43]. Considerable genetic and transcriptomic variability have been shown among different *Pv* strains [14, 44, 45], but how such variation influences survival and development of the parasites between *in vivo* and *ex vivo* environments remain unclear.

To address these knowledge gaps, this study investigated paired *in vivo* and short-term *ex vivo Pv* transcriptomes from malaria patients in Ethiopia, with the goals to (1) compare inferred developmental stage composition between conditions; (2) determine whether short-term *ex vivo* culture alters parasite transcriptomes after accounting for differences in parasite stage; (3) examine within-host clonal dynamics between *in vivo* and *ex vivo* samples using amplicon deep sequencing and; (4) explore whether residual transcriptional heterogeneity in low-complexity infections is associated with dominant haplotype. These comparisons provide a framework for interpreting *Pv* transcriptomes in the context of developmental asynchrony and within-host genetic diversity.

## Results

### Short-term *ex vivo* culture reshapes developmental stage composition

We obtained high-quality transcriptomes from 40 paired *Pv* clinical isolates processed directly from patient blood (denoted as *in vivo* hereafter) and after one intraerythrocytic cycle under short-term culture conditions (denoted as *ex vivo*; S1 Table). We first assessed whether the *ex vivo* condition altered parasite developmental composition profiles. Gene expression deconvolution analysis indicated clear differences in parasite stage distributions among samples and conditions. Across both conditions, trophozoites comprised the dominant inferred fraction (median *in vivo* 0.532 vs *ex vivo* 0.530; Wilcoxon *p*=0.816; Table 1), indicating overall preservation of the major asexual transcriptional signal. However, paired analyses revealed significant changes in composition with a relative enrichment of late asexual stages following culture (Fig 1A-B; Table 1). Across 40 matched *in vivo*–*ex vivo* pairs, the proportion of mRNA derived from schizonts significantly increased after *ex vivo* culture (median 0.027 vs 0.097; *p*=0.008), consistent with continued maturation of circulating parasites during the culture interval (Fig 1C; Table 1). In particular, in many samples, schizont mRNAs were undetectable or very low in vivo but significantly increased after ex vivo culture. In contrast, the proportion of ring-stage parasites decreased after culture (median 0.174 vs 0.092; *p*=0.056; Fig 1D). Sexual-stage gametocyte fractions were generally similar. Female gametocyte (median 0.166 vs 0.133; *p*=0.563) and male gametocyte fractions (median 0.040 vs 0.040; *p*=0.254) did not differ significantly between conditions (Fig 1E-G), although there were substantial inter-isolate variations. It is worth noting that several infections showed high inferred female gametocyte fractions at clinical presentation (Fig 1G), implicating the frequent presence of transmissible stages in symptomatic infections. Collectively, these data indicate that even not a full short-term culture cycle modestly reshaped developmental stage composition, enriching for late asexual stages while preserving trophozoite and gametocyte signals in the transcriptome.

**Figure 1.**
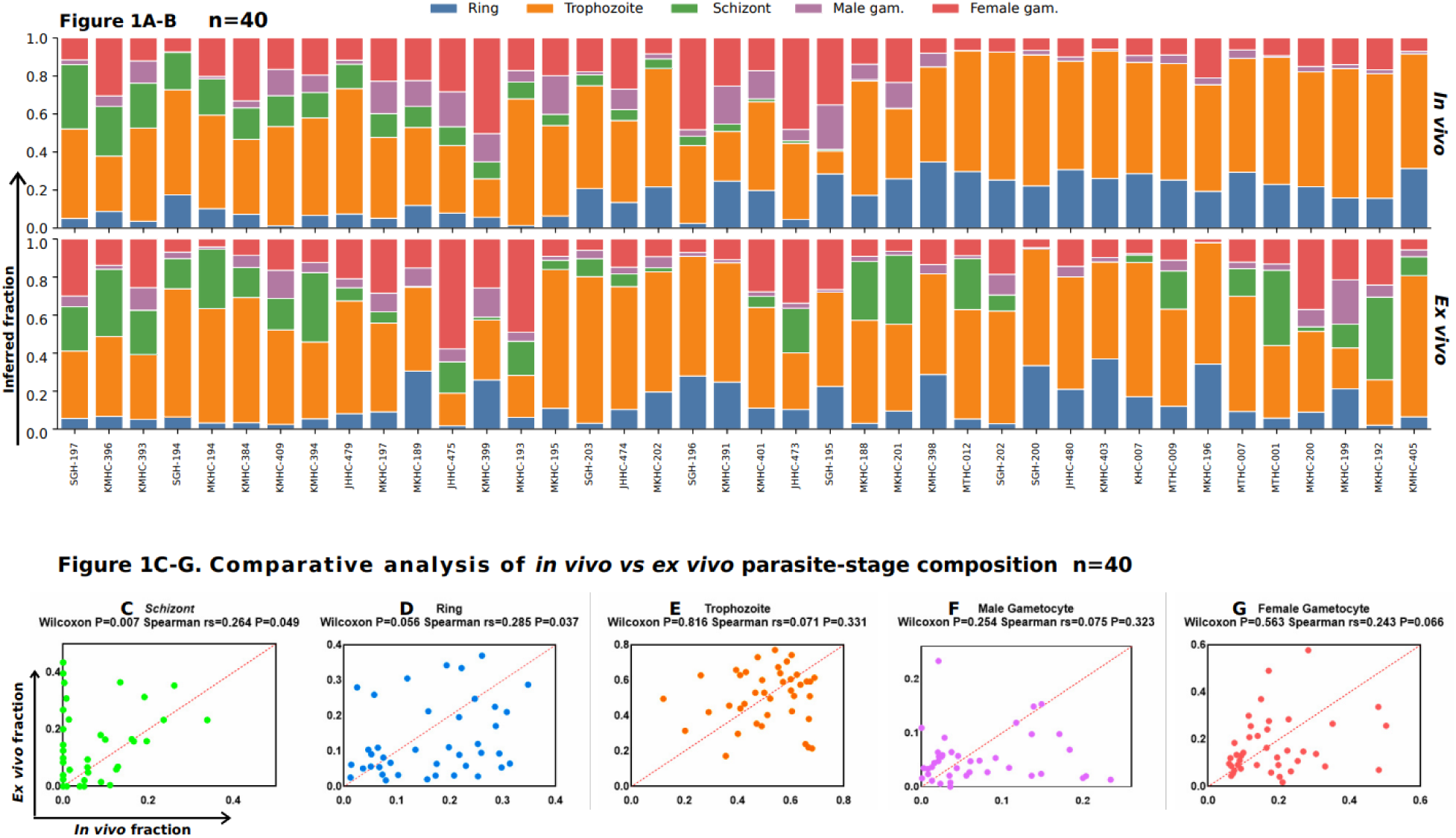
Short-term *ex vivo* culture alters parasite-stage composition in paired *Pv* clinical isolates. **(A-B)** Stacked bar plots showing inferred developmental stage composition for 40 matched clinical isolates processed directly from patient blood (*in vivo*, top panel) and after one intraerythrocytic cycle under short-term culture conditions (*ex vivo*, bottom panel). Each bar represents one isolate, and the colors denote developmental stages: ring (blue), trophozoite (orange), schizont (green), male gametocyte (purple), and female gametocyte (red). **(C-G)** Paired comparative analysis of inferred stage fractions between *in vivo* and *ex vivo* conditions. Each point represents a paired isolate (*n*=40) displayed according to the proportion of a given stage in vivo (x-axis) and ex vivo (y-axis). The dashed red line indicates the line of identity (y=x). Wilcoxon signed-rank tests were used for paired comparisons; Spearman correlation coefficients (rs) are shown to illustrate rank concordance between conditions.

**Table 1.**
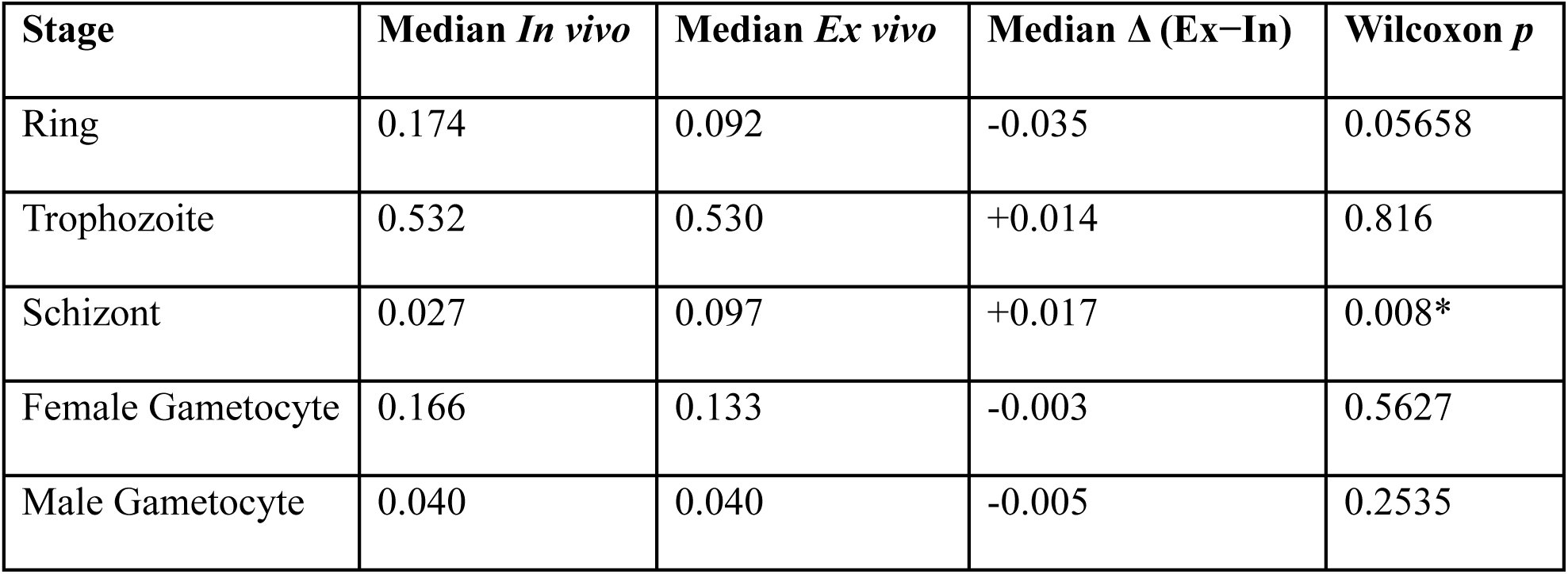
Paired comparison of stage fractions (*ex vivo* vs in vivo; n = 40 pairs). Asterisk denotes significant difference at *p*=0.05.

| Stage | Median *In vivo* | Median *Ex vivo* | Median $\Delta$ (Ex–In) | Wilcoxon *p* |
| --- | --- | --- | --- | --- |
| Ring | 0.174 | 0.092 | -0.035 | 0.05658 |
| Trophozoite | 0.532 | 0.530 | +0.014 | 0.816 |
| Schizont | 0.027 | 0.097 | +0.017 | 0.008* |
| Female Gametocyte | 0.166 | 0.133 | -0.003 | 0.5627 |
| Male Gametocyte | 0.040 | 0.040 | -0.005 | 0.2535 |

### No significant difference in stage-adjusted transcription between *in vivo* and *ex vivo* conditions

To test whether short-term *ex vivo* culture induces a transcriptional program beyond developmental redistribution, we performed both uncontrolled and stage-adjusted analyses of paired *in vivo* and *ex vivo* transcriptomes. PCA showed extensive overlap between *in vivo* and *ex vivo* samples (Fig 2A), with no clear separation observed along the major axes of variation (PC1=33.3%, PC2=11.0%), indicating that culture condition does not dramatically impact the expression profile of the parasites. Consistent with this observation, stage-adjusted paired differential gene expression (DGE) comparisons showed no genes significantly differentially expressed at FDR=0.05 (Fig 2B and S2A Fig). The volcano plot revealed a symmetric distribution, with no genes reaching statistical significance after multiple-testing correction. Only a small number of genes reached FDR<0.10, and these residual signals after stage correction were modest in magnitude and dominated by variant/exportome-like annotations, including several PIR family members (Table 2).

**Figure 2.**
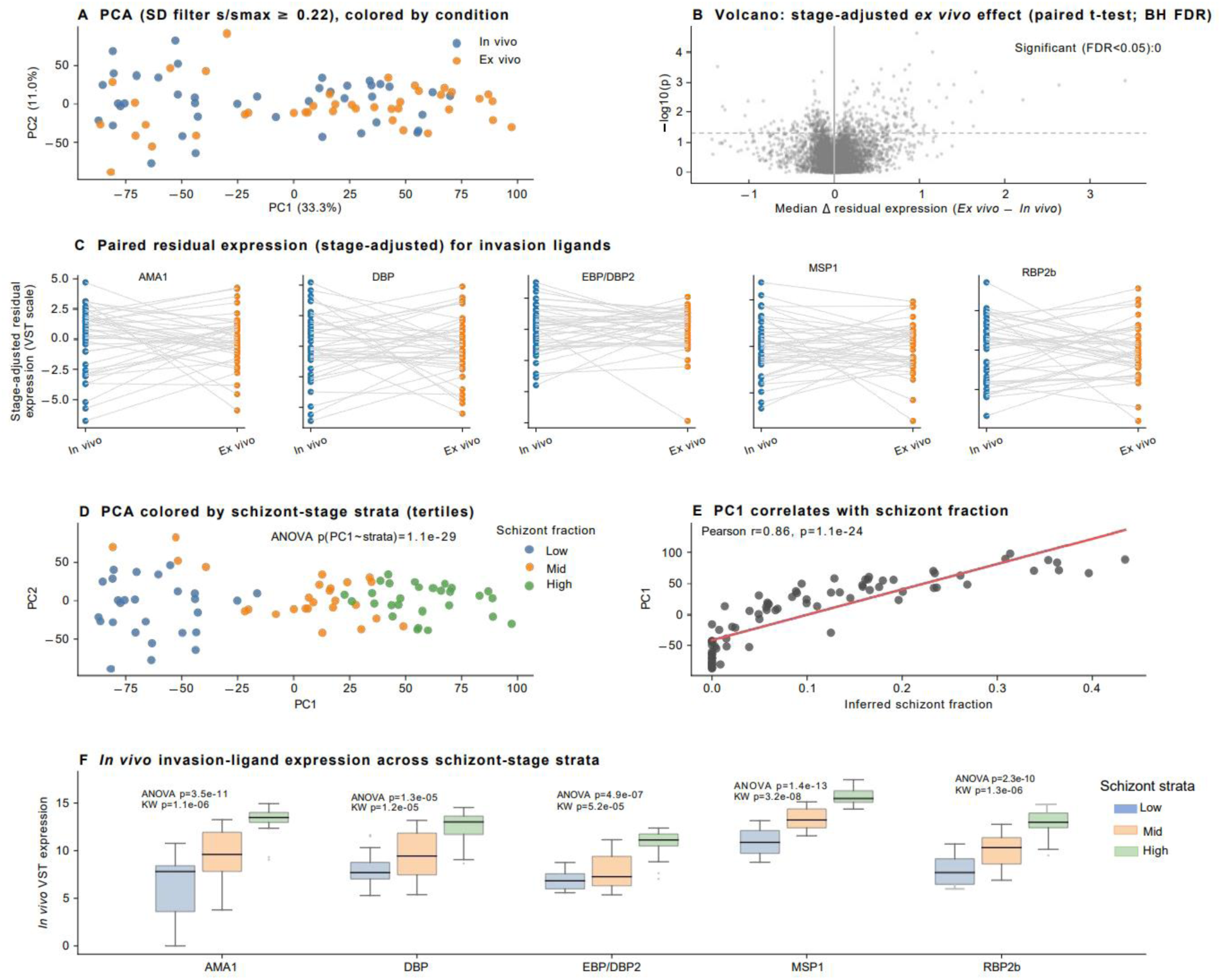
Stage composition determines transcriptome structure, whereas stage-adjusted *ex vivo* condition effects were minimal. **(A)** Principal component analysis (PCA) of variance-stabilized parasite gene expression values colored by condition (*in vivo* vs *ex vivo*). Samples showed extensive overlap with no clear separation by culture exposure. **(B)** Volcano plot of stage-adjusted differential expression for the *ex vivo* effect (paired t-test with Benjamini-Hochberg correction). The x-axis showed median fold change (*ex vivo* - *in vivo*), and the y-axis shows - log10(p). No genes met significance at FDR<0.05. The distribution of fold changes was symmetric around zero. **(C)** Paired stage-adjusted residual expression (variance-stabilized scale) for key invasion and merozoite-associated genes. Each line represents one isolate in vivo (left) and ex vivo (right). No systematic residual shift was observed between *in vivo* and *ex vivo* samples. **(D)** PCA colored by inferred schizont-stage tertiles (low, mid, high). PC1 stratified strongly by schizont fraction (ANOVA *p*=1.1 × 10⁻²⁹), indicating that developmental composition, rather than culture condition, dominates transcriptome organization. **(E)** Correlation between PC1 and inferred schizont fraction. PC1 correlated strongly with schizont abundance (Pearson *r*=0.86, *p*=1.1 × 10⁻²⁴), demonstrating that late asexual-stage representation is a major determinant of overall transcriptional variance. **(F)** *In vivo* invasion-ligand expression across schizont-stage tertiles. Variance-stabilized expression of invasion and merozoite associated markers increased significantly across increasing schizont proportions (ANOVA and Kruskal-Wallis *p*<0.001 for all), consistent with their expression at this stage.

**Table 2.** Top stage-adjusted signals (FDR < 0.10).

| Gene ID | Log2FC(adj) | FDR | Gene function |
| --- | --- | --- | --- |
| PVP01_0121700 | +1.15 | 0.080 | PIR protein |
| PVP01_0534700 | +0.63 | 0.080 | PIR protein |
| PVP01_0009800 | +0.90 | 0.080 | PIR protein |
| PVP01_0004380 | +0.70 | 0.080 | PIR protein |
| PVP01_1461400 | +0.65 | 0.080 | ACEA small nucleolar RNA U3 |

Examination of key invasion and merozoite associated genes, including *PvDBP*1, *PvEBP/DBP*2, *PvRBP*2b, *PvMSP*1, *PvAMA*1, similarly showed no consistent stage-adjusted differences between paired *in vivo* and *ex vivo* samples (Fig 2C; paired *t*-test *p*>0.05). In contrast, the overall transcriptome structure was strongly organized by developmental state. For instance, along PC1 clear differences were observed in the transcriptome signal when samples were stratified by inferred schizont fraction (ANOVA *p*=1.1×10⁻²⁹; Fig 2D). PC1 correlated strongly with inferred schizont fraction (Pearson *r*=0.86, *p*=1.1×10⁻²⁴; Fig 2E), demonstrating that late asexual-stage representation was the dominant determinant of transcriptomic variance. Within *in vivo* infections, invasion-ligand expression increased significantly across schizont tertiles for multiple invasion- and merozoite-associated markers (Fig 2F), consistent with stage-dependent regulation of invasion gene expression. These findings suggested that the apparent “*ex vivo* effects” on late-stage genes are largely explained by shifts in developmental composition. After controlling for stage, short-term *ex vivo* culture did not induce an overall similar transcriptional signal across isolates. Because direct stage-adjusted condition effects were minimal, we next examined whether residual transcriptional signals were associated with within-host parasite genotypes.

### Short-term culture imposes a clonal bottleneck and non-random selection among *PvMSP*1 haplotypes

The highly variable Block 2 region of the *P. falciparum* or *P. vivax* MSP1 gene is commonly used as a proxy for genetic diversity. To quantify within-host clonal diversity and directly track genotype dynamics across conditions, we performed *PvMSP*1-based haplotyping in 28 paired *in vivo* and *ex vivo* samples. Haplotypes were defined by read-based detection thresholds (presence ≥5% of reads), allowing determination of multiplicity of infection (MOI), identification of dominant haplotypes, and tracking of clone persistence across the paired samples. A total of 31 unique *PvMSP*1 haplotypes were detected across the dataset, of which 28 were observed in *in vivo* and 28 *in ex vivo* samples (S3 data). Short-term ex vivo culture altered parasite clonal composition: *ex vivo* samples showed a marked shift towards lower complexity infections, with a reduction in highly polyclonal (>3 clones) infections and an increase in monoclonal and biclonal states, consistent with a contraction of within-host diversity during *ex vivo* condition (Fig 3A). Despite this reduction, several haplotypes persisted across paired samples, most frequently H-4 (detected in 11 pairs) and H-7 (9 pairs), followed by H-17, H-1, and H-10 (5 pairs each; S3 data). However, haplotype prevalence differed between conditions in a subset of infections. In six of the 28 matched pairs, haplotypes were not shared between *in vivo* and *ex vivo* samples, indicating clone loss or emergence (i.e. above detection level) when the condition changes. Fisher’s exact enrichment analysis identified a single haplotype, H-29, to be significantly more abundant in *in vivo* (17/28, 58.6%) relative to *ex vivo* samples (2/28, 6.9%; OR*=*19.125, 95% CI 3.803-96.184, *p*=4.7×10⁻⁵, q=0.001; Fig 3B). The odds-ratio distribution (log scale) with values >1 indicating *in vivo* enrichment (blue) and <1 indicating *ex vivo* enrichment (red), with only H-29 exceeding the FDR significance (q<0.05). For dominant haplotypes (e.g., H-4, H-7, H-10), mean relative abundance was broadly similar between the two conditions (Fig 3B), although modest shifts were observed for some haplotypes, such as H-29 that exhibited marked reduction in *ex vivo* relative to *in vivo* conditions (Fig 3B; S3 data). To quantify directional changes in clonal composition, we examined the net haplotype change within each matched infection. Most pairs exhibited a reduction in haplotype complexity following short-term culture, with losses of two (n=12), one (n=6), or three (n=3) haplotypes being the most common outcomes (Fig 3C). Complete stability (n=2) and gains in haplotype number (gain of one, n=4; gain of three, n=1; Fig 3C) were less common. These data demonstrated an overall contraction in within-host diversity. Importantly, this contraction was not random. Instead, specific haplotypes displayed consistent directional shifts in frequency across samples, with some enriched while others depleted.

**Figure 3.**
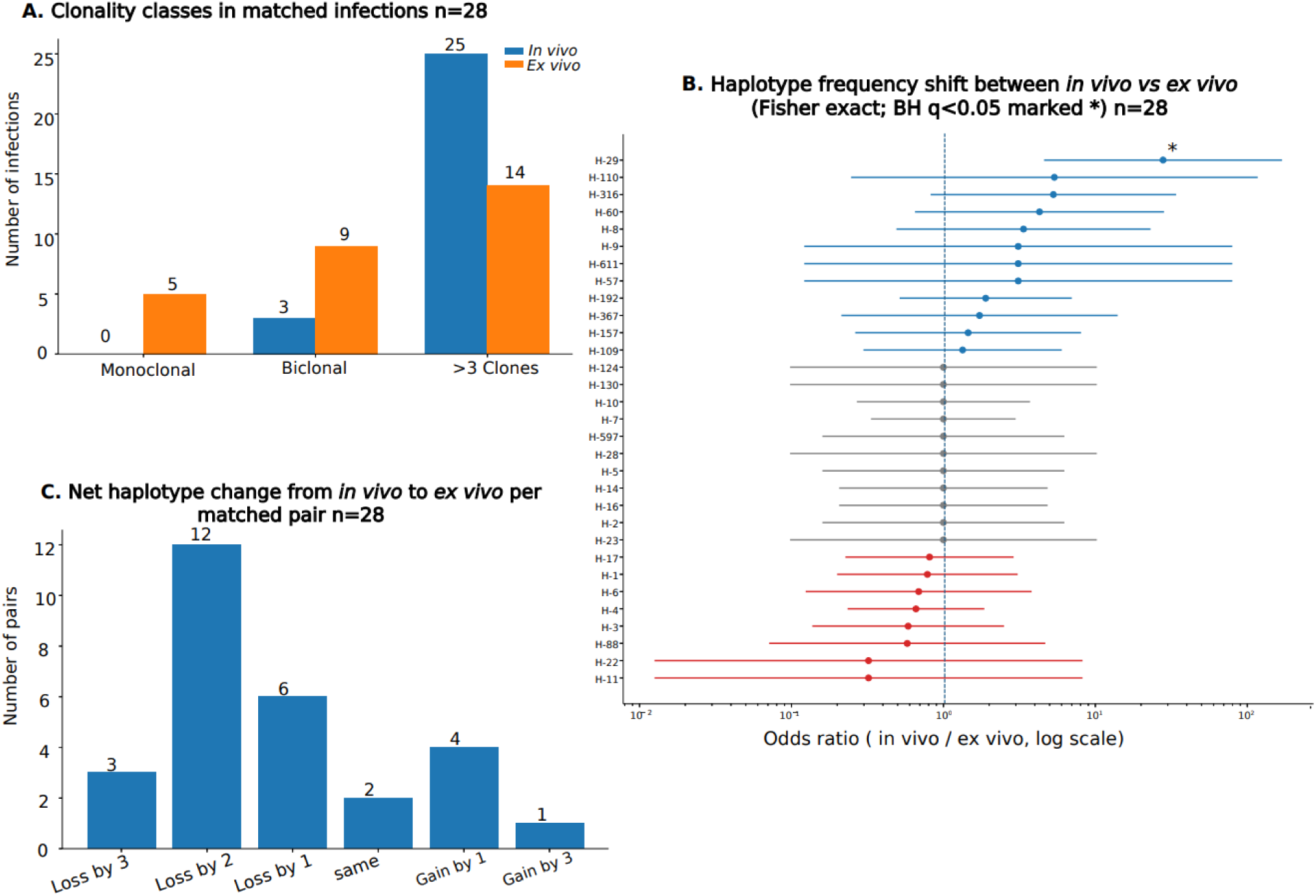
Short-term *ex vivo* culture decreases clonal diversity and non-random selection of *PvMSP*1 haplotypes. **(A)** Distribution of multiplicity of infection (MOI) categories across 28 paired samples *in vivo* (blue) and after short-term *ex vivo* culture (orange), classified as monoclonal, biclonal, or >3 clones based on *PvMSP*1 haplotypes detected at ≥5% read frequency. *Ex vivo* samples show a shift towards lower complexity infections, with reduced representation of highly polyclonal (>3 clones) infections and increased monoclonal and biclonal states, consistent with a contraction of within-host diversity during ex vivo culture. **(B)** Haplotype-specific frequency shifts between *in vivo* and *ex vivo* conditions. Forest plot of odds ratios (*in vivo*/*ex vivo*, log scale) for each *PvMSP*1 haplotype across 28 paired infections, estimated using Fisher’s exact test with Benjamini–Hochberg correction. The vertical dashed line indicates no difference (OR=1). Haplotypes with OR>1 (blue) were enriched *in vivo*, whereas those with OR<1 (red) were enriched *ex vivo*. Error bars represent 95% confidence intervals. Only haplotype H-29 showed significant enrichment *in vivo* after multiple testing correction (q<0.05; marked with *), indicating consistent depletion during short-term culture and supporting non-random selection among parasite clones. **(C)** Distribution of the number of haplotypes lost or gained between *in vivo* and *ex vivo* samples for each of the 28 matched pairs. Most infections exhibited a reduction in haplotype number, with losses of two (n=12), one (n=6), or three (n=3) haplotypes being the most common outcomes, whereas complete stability (n=2) and haplotype gains (gain of one, n=4; gain of three, n=1) were less common. This asymmetric pattern indicates a directional bottleneck effect rather than stochastic turnover of parasite haplotypes.

Although the paired dataset (Table 3; *n*=28) showed a significant reduction in median MOI, from 4.0 *in vivo* to 2.5 *ex vivo* (median paired change −1.0; Wilcoxon *p*=0.001; Table 3), 78.6% of all haplotypes detected *ex vivo* were present in the corresponding *in vivo* sample, yielding an overall haplotype survival rate of 0.609 (Table 3). The probability of a dominant haplotype persisted after culture increased with its initial abundance, with each 0.10 increase in *in vivo* dominant haplotype fraction associated with higher odds of persistence (OR=2.87, *p*=0.012; Table 3). This analysis further indicated that clone persistence was associated with higher initial abundance, rather than stochastic survival.

**Table 3.** summary of MSP1 microhaplotype-based clonal metrics (paired samples: n=28).

| Metrics | Value |
| --- | --- |
| Paired patients with MSP1 data | N=28 (*in vivo* and *ex vivo* paired) |
| Haplotype persists | 22/28 (79%) |
| Haplotype switches | 6/28 (21%) |
| Median MOI (*In vivo*) | 4.0 |
| Median MOI (*ex vivo*) | 2.5 |
| Median MOI change (*ex vivo* – *in vivo*) | -1.0 |
| Paired MOI analysis across all haplotypes Wilcoxon p | 0.001 |
| Haplotype survival rate (*in vivo* → *ex vivo*) | 0.609 |
| *Ex vivo* haplotypes pre-existing *in vivo* | 0.786 |
| Persistence OR per =0.10 *in vivo* dominant fraction | 2.87 (*p*=0.012) |

### Exploratory dominant haplotype-associated residual expression patterns in low-complexity subset samples

To examine whether parasite genotype contributes to transcriptional heterogeneity beyond developmental stage effects and culture condition, we analyzed variance-stabilized gene expression values after adjustment for parasite stage, condition, and parasitemia. Analyses were restricted to low-complexity infections (MOI ≤2; n=12) to minimize confounding signals from within-host clonal mixtures. Principal component analysis (PCA) of the adjusted expression matrix revealed distinct clustering of samples according to the dominant *PvMSP*1 haplotypes (H-4, H-7, and H-17) (Fig 4A). This clustering pattern persisted despite explicit control for parasite stage composition, indicating that residual transcriptional variation was not driven by developmental differences but instead reflects genotype-associated expression programs. To quantify the genotype-associated effect, we performed a multi-group ANOVA on the adjusted expression matrix. While a significant nominal association was detected between dominant haplotype and gene expression (R²=0.70; *p*=0.004), gene-level differences did not remain significant after multiple testing correction (q=0.686). Thus, these results suggested an exploratory rather than definite haplotype-associated transcriptional effect.

**Figure 4.**
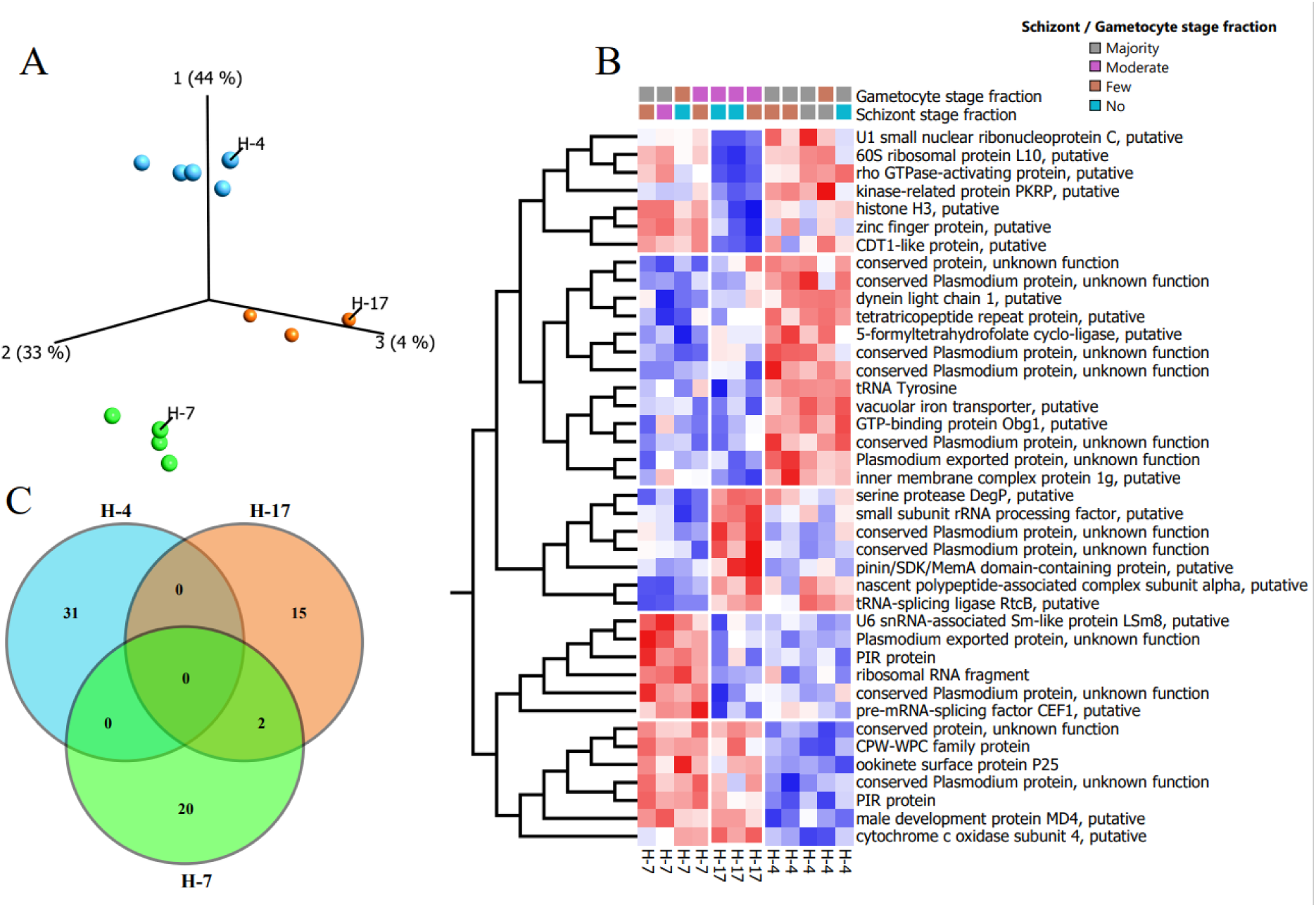
Dominant haplotype exhibited a distinct stage-adjusted transcriptional program. **(A)** Principal component analysis (PCA) of samples by dominant *PvMSP*1 haplotype (H-4, H-7, H-17) among low complexity infections (MOI ≤2; *n* = 12), illustrating distinct clustering of haplotypes. Percent variance explained by each principal component was indicated. **(B)** Hierarchically clustered heatmap using multi-group ANOVA performed on the residual, standardized expression matrix demonstrated a significant nominal association between dominant haplotype and gene expression (R²=0.70; *p*=0.004), although this effect did not remain significant after multiple-testing correction (*q*=0.686). Rows represent genes and columns represent samples, ordered by dominant haplotype. Expression values are standardized Z-score per gene. Dominant haplotype samples exhibit a coherent expression block distinct from other haplotypes. Top annotations indicated samples stratified by schizont and gametocyte fractions, demonstrating that clustering is independent of stage composition. **(C)** Venn diagram showing the overlap of genes among haplotypes. Several genes were uniquely associated with individual haplotype, including 31 genes unique to H-4, 20 genes unique to H-7, and 15 genes unique to H-17 with limited overlap, indicating haplotype-specific transcriptional programs.

Inspection of the heatmap revealed coherent gene modules that varied systematically across haplotypes (Fig 4B). H-4 dominated infections exhibited elevated expression of genes associated with translation and cellular and biosynthetic processes, including ribosomal proteins (e.g., 60S ribosomal protein L10), RNA processing factors (e.g., U1 small nuclear ribonucleoprotein C, LSm proteins), and chromatin-associated proteins such as histone H3. This module suggested a metabolically active, proliferative state associated with asexual replication and invasion. In contrast, H-7 dominated infections showed increased expression of PIR family members and exported proteins (e.g., PVP01_0000570, PVP01_1000300), indicative of immune evasion and antigenic variation. The upregulation of these genes suggested that H-7 parasites may preferentially engage host-interaction pathways, potentially reflecting an adaptive strategy focused on persistence within the host environment. H-7 and H-17 dominated infections were characterized by increased expression of genes associated with sexual differentiation and gametocytes, including P25 (okinete surface protein PVP01_0616100), CPW-WPC family proteins, as well as male development-associated factors (e.g., MD4). This module was consistent with a transmission-biased transcriptional state, suggesting that H-7 and −17 dominated infections may be skewed towards gametocyte development. Pairwise comparisons further supported haplotype-specific programs, with the strongest contrasts observed between H-4 and H-17, and between H-4 and H-7, across multiple genes. A Venn diagram of differentially expressed genes between haplotype pairs demonstrated that most transcriptional differences were haplotype-specific (Fig 4C), including 31 genes unique to H-4, 20 genes unique to H-7, and 15 genes unique to H-17, with minimal overlap. Only two genes were shared between H-7 and H-17, indicating that each haplotype was characterized by a distinct transcriptional program. These results illustrated that dominant *PvMSP*1 haplotypes had distinct residual transcriptional states after controlling for stage composition, culture condition, and parasite load.

## Discussion

By integrating paired *in vivo* and short-term *ex vivo* transcriptomes with developmental-stage deconvolution and *PvMSP*1 haplotyping, this study shows that much of the transcriptional differences between *in vivo* and *ex vivo Pv* parasites were explained by changes in developmental stage composition rather than a shared culture-induced transcriptional response. Short-term *ex vivo* culture modestly altered developmental composition, primarily enriching late asexual (schizont) stages. After accounting for stage composition, the overall expression profiles of *in vivo* and *ex vivo* isolates remained similar, with no genes differentially expressed at FDR<0.05. In a small subset of low-complexity infections, residual expression patterns were associated with dominant haplotype, indicative of genotype-linked heterogeneity that warrants further validation. These findings suggested that developmental asynchrony and polyclonality are not merely confounders [6, 7, 22, 31, 32, 46–48] but central biological features shaping the *Pv* transcriptome.

The developmental shifts observed after one erythrocytic cycle *ex vivo* were biologically expected and help reconcile discrepancies between transcriptomes obtained directly from patient blood and those derived from short-term maturation [6, 22, 26, 27, 29, 46]. Blood-stage *Pv* transcription profiles are strongly influenced by the parasite developmental stage, and late asexual stages are expected to disproportionately contribute signal from merozoite- and invasion-related genes [2, 7, 23, 26, 29]. Consistent with this framework, our data showed that *ex vivo* culture enriched schizont-stage signal, whereas trophozoite and gametocyte fractions were largely preserved. The strong correlation between transcriptomic variance (PC1) and inferred schizont fraction further supports the notion that stage composition, rather than ex vivo culture, is the dominant determinant of global transcriptional structure. Accordingly, the stronger invasion-gene signal in *ex vivo* samples is best interpreted as a consequence of stage redistribution rather than induction of a distinct culture-specific transcriptional state. These finding are consistent with single-cell atlases of *Pv* blood stages and prior bulk transcriptomic studies, which showed that stage composition is a primary determinant of transcriptional variation across infections [7, 23, 47]. The broad overlap observed between paired *in vivo* and *ex vivo* samples after stage adjustment also aligns with earlier conserved blood-stage transcriptional program despite isolate-specific variability and complex isoform usage [29, 46]. This has an important practical implication because in the absence of a continuous *Pv* culture system, short-term *ex vivo* maturation remains one of the few feasible approaches for enriching late asexual stages [1, 4, 5, 48]. Our results indicated that such approaches could yield biologically meaningful data, provided that developmental composition is explicitly modeled prior to interpretation [6, 22, 26, 47].

Despite the absence of detectable effect of ex vivo culture on the parasite gene expression, we observed a consistent reduction in within-host clonal diversity following short-term *ex vivo* maturation, this is consistent with the difficulty to propagate *Pv* in vitro condition. *PvMSP*1 haplotyping showed a decrease in multiplicity of infection and frequent loss of one or more haplotypes per infection, consistent with a clonal bottleneck. Polyclonal and monoclona *Pv* infections are common in endemic settings and can arise through repeated exposure, superinfection, and relapse from multiple dormant liver-stage parasites [31–37, 49, 50]. In this study, the non-random reduction in MOI *ex vivo*, the persistence of dominant haplotypes (e.g., H-4, H-7, and H-17), the dependence of clonal persistence by initial abundance and stage, and the selective depletion of specific haplotypes (e.g. H-29) together support genotype-dependent selection rather than stochastic dropout. These findings suggest that short-term culture preferentially retains clones with greater *ex vivo* fitness, potentially reflecting variations in growth rate, maturation efficiency, invasion capability, or tolerance of altered host-cell conditions among individual clones [14, 26, 46, 51]. Genotype-associated transcriptional structure further supports this interpretation. In stage-, condition-, parasitemia-adjusted expression profiles, dominant *PvMSP*1 haplotypes were associated with residual expression signals in a subset of low-complexity infections. In particular, H-4 dominated infections showed clear clustering and were associated with gene modules linked to translation, cellular growth, and chromatin regulation, whereas H-7 and H-17-dominated infections showed stronger PIR/exported-protein signatures and gametocyte features, respectively. Although limited by sample size, these data suggest that co-circulating parasite clones may occupy distinct functional states. Extensive diversity in invasion ligands and host-interaction genes could make genotype-linked expression biologically plausible and may contribute to differential survival under *ex vivo* conditions. Larger datasets will be required to validate the genotype-associated transcription signals. In *P. falciparum*, within-host genetic complexity has been linked to altered expression patterns, longer gametocyte carriage, and host-mediated selection among co-circulating genotypes [38–41]. The observed genotype-associated residual expression in *Pv* and prior observations from *P. falciparum* indicated that parasite genetic background can organize transcriptional variation independently of stage. It is plausible that variation in clonal fitness and regulatory state may already exist within infections at the time of clinical presentation and become more apparent under *ex vivo* conditions that impose selective pressure.

Taken together, these results support a model in which short-term *ex vivo* culture primarily filters pre-existing within-host diversity rather than inducing a uniform transcriptional response. During *ex vivo* maturation, predictable enrichment of late asexual stages is superimposed on non-random changes in clonal representation. This framework explains why paired *in vivo* and *ex vivo* transcriptomes remain globally similar after stage adjustment while still showing pronounced turnover in haplotype composition and genotype-associated residual expression. These findings have several implications for future *Pv* transcriptomic studies. First, comparison between *in vivo* and *ex vivo* samples must account for stage composition before attributing biological difference to culture exposure. Second, while short-term *ex vivo* maturation is a useful tool for enriching late stages, it may selectively favor specific clones and therefore not fully represent the within-host diversity present at clinical presentation. Third, studies aiming to identify invasion pathways, transmission biology, or isolate-specific features from bulk RNA-seq should explicitly incorporate both stage deconvolution and measures of infection complexity. More broadly, our results emphasize that asynchronous development and polyclonality are intrinsic properties of natural *Pv* infections that shape the observed transcriptome that must be considered in analyses.

Despite these important implications, there are few limitations in this study. First, bulk RNA-seq, even when corrected for stage composition, does not reflect signals of minor clones or rare developmental states in highly asynchronous infections. Second, *PvMSP*1 haplotypes are useful for tracking clonal dynamics but do not resolve whole-genome haplotypes. Third, haplotype-stratified analyses relied on modest numbers of low-complexity infections, limiting the power for less common haplotypes. Finally, because this study used bulk transcriptomes from one endemic setting, the generalizability of clone-specific transcriptional signals remains to be tested in other geographical isolates. Recent advances in single-cell preservation, field-compatible scRNA-seq, and analysis of polyclonal *Pv* infections should help address these limitations in future work by directly resolving clone- and stage-specific programs within mixed infections.

## Conclusion

This study provides a framework for interpreting *Pv* transcriptomes in the context of two fundamental features of natural infections: developmental asynchrony and polyclonality. After explicit adjustment for parasite stage, short-term *ex vivo* culture did not induce a shared transcriptional program. Instead, the most informative residual variation reflected non-random clonal survival and dominant haplotype-associated expression signals. These findings suggest that *Pv* infections are not homogeneous parasite populations, but dynamic assemblages of genetically distinct lineages with different functional states. Short-term *ex vivo* culture primarily acts as a selective filter, enriching late asexual stages while imposing a clonal bottleneck that favors specific parasite genotypes. This process reflects genotype-dependent fitness differences rather than stochastic loss of diversity and results in measurable shifts in clonal composition without global transcriptional reprogramming. Collectively, our findings highlight the need to explicit model both developmental composition and clonal complexity when analyzing field-isolate transcriptomes. Doing so will be essential for accurately identifying parasite biological features relevant to natural infections and for improving the design and interpretation of *ex vivo* functional studies.

## Materials and Methods

### Ethical statement

The study protocol was reviewed and approved by the National Research and Ethics Review Committee (NRERC) of Jimma University, Ethiopia (Ref. No.03/246/796/22) and Drexel University, USA. Written informed consent/assent was obtained from all consenting heads of household, parents/guardians, and individuals who were willing to participate in the study. All experimental procedures were performed following the IRB approved protocol.

### Sample collection and processing

Blood samples of 45 *Pv*-infected individuals from southwestern Ethiopia were collected between March-April 2024. These samples were microscopy and RDT confirmed *Pv* mono-infection and absence of antimalarial treatment in the preceding 30 days. Ten milliliters of venous blood were collected into heparin tubes. Plasma was isolated within 1 hour, and the red blood cell pellet was divided and preserved in two ways: (1) *in vivo*: 5 mL packed cells in 10 volumes of TRIzol™ (Thermo Fisher) and stored at −80°C overnight followed by liquid nitrogen storage; (2) *ex vivo*: the remaining pellet was washed in incomplete IMDM medium and cultured in 35 mL complete IMDM supplemented with (2.5% human AB plasma, 2.5% HEPES buffer, 2% hypoxanthine, 0.25% albumax, and 0.2% gentamycin) for 24-48 h at 37°C until the majority of parasites reached the schizont stage with a final ∼2% hematocrit inside a CO2 controlled incubator with <5% O2 level. Cultured cells were pelleted by centrifugation and preserved in TRIzol™. To control differences in parasite stage, parasite growth was monitored by microscopy of a thick smear prepared from the culture every 4, 8, and 16 hours after the initial starting time, depending on the majority stage. To minimize oxidative stress, replicate cultures were monitored microscopically and returned immediately to a 5% oxygen environment after each assessment. When cultures reached >20% mature schizonts ex vivo, samples were centrifuged and pellets containing both infected and uninfected RBC were placed in 10×trizol.

### RNA extraction, library preparation, and sequencing

Total RNA and DNA were extracted simultaneously from TRIzol™-preserved pellets using phenol-chloroform phase separation followed by isopropanol precipitation. Eluted in a final volume of 60µl. RNA quality was assessed by Qubit™ 4 Fluorometer and Agilent Fragment Analyzer; samples with RNA Integrity Number (RIN≥6, DV200≥70%, and concentrations exceeded 150 ng/μL were selected for library construction. Ribosomal RNA and globin was depleted using the NEBNext® rRNA and globin Depletion Kit (Human/Mouse/Rat), followed by strand-specific library preparation with the NEBNext® Ultra™ II Directional RNA Library Prep Kit. Libraries were quantified by Qubit™ and quality-checked on an Agilent Bioanalyzer. Paired-end sequencing (2×150 bp) was performed on the Illumina NovaSeq 6000 platform at Maryland Genomics to generate ≥ 35 million reads per sample. Across all libraries, sequencing yielded a mean of 120,504,651 reads per sample (range 87,322,339-189,622,359). After alignment, a mean of 51,001,345 reads mapped to the parasite reference (mean 40%, range 5-75%), consistent with variable parasite biomass and host background across clinical specimens.

### RNA sequencing and data processing

Adapter sequences and low-quality bases were removed using Trimmomatic v0.39. To confirm single *Plasmodium* species infection, all patient samples were mapped using HISAT2 v2.1.0 [52] against a fasta file containing the genome, filtered out for mitochondrial sequences, of the 5 human infecting *Plasmodium* species: *P. vivax* P01, *P. falciparum* 3d7, *P. knowlesi* GCF, *P. malariae* UG01, and *P. ovale curtisi* GH01 (PlasmoDB [53, 54], v68). Finally, all samples were realigned to *P. vivax* P01genome with a maximum intron length of 5,000 bp, uniquely mapping reads were parsed using samtools v1.20 [55], and PCR duplicates were removed using Picard MarkDuplicates v2.25.3 (Broad Institute). Reads were further assigned to specific protein-coding gene and counted using the subread feature Counts v2.0.6 [56] based on the *P. vivax* genome annotation PlasmoDB (Release 68, May 7, 2024).

### Gene expression analysis

Raw counts were used for count-based differential expression modeling. Read counts were normalized via VST for differential expression analyses. Statistical assessment of differential expression was conducted, for the *in vivo* and *ex vivo* samples to assess the *ex vivo* effect*, in DESeq2* with and without adjusting for proportion of each parasite developmental stage for *Plasmodium vivax* genes[57, 58]. Relative proportions of rings, trophozoites, schizonts, male gametocytes, and female gametocytes were inferred using CIBERSORTx [47, 59, 60]. For exploratory dominant-haplotype analyses, gene expression values were adjusted for parasite stage composition and experimental covariates using gene-wise linear regression. For each gene independently, we fit the following model:

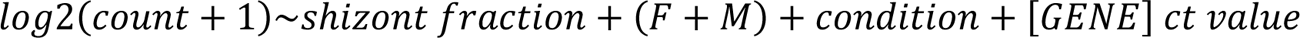

where schizont and gametocyte fractions were obtained from deconvolution estimates from CIBERSORTx, condition was encoded as a binary variable (*in vivo*=0, *ex vivo*=1), and Ct values targeting 18S rDNA gene of the parasite and based on using established protocol[61, 62] utilizing species-specific primers (forward: 5’-AGTCATCTTTCGAGGTGACTTTTAGATTGCT-3’; reverse: 5’ GCCGCAAGCTCCACGCCTGGTGGTGTC-3’) were used as a proxy for parasite density. Residuals from these models were extracted and used as stage- and condition-adjusted expression values.

To enable cross-gene comparability and visualization, residualized expression values were further normalized by computing a per-gene Z-score across samples (mean=0, variance=1). To ensure non-negative values suitable for downstream visualization Z-scored values were shifted such that the minimum value for each gene was set to zero. Analyses were performed on complete-case samples with available covariate information; samples with missing covariates were excluded from regression but retained as missing values in the full matrix outputs.

### *PvMSP*1 Genotyping for Clonal Identity

Clonal identity was assessed by genotyping the polymorphic *P. vivax merozoite surface protein-1* (*PvMSP1*) gene. *In vivo PvMSP*1 genotyping was performed on DBS-derived DNA. Briefly, Genomic DNA was extracted from *Pv* clinical isolates using a standard saponin/Chelex method as previously described Bereczky *et al.*, 2005. Whereas *ex vivo* MSP1 genotyping was performed on DNA recovered from matched cultured TRIzol-preserved pellets. We employed a two-step PCR amplicon sequencing protocol adapted from a published method. In the first PCR, ∼463 bp of the highly variable region of *PvMSP1* (PVX_099980, NCBI reference AF435593) was amplified using gene-specific primers (Forward primer: 5’ ACCCATACAAGCTGCTCGAC 3’ and Reverse primer: 5’ TCCTCCAACTTCTCATCCATC 3’) with Illumina adapter tails (**Fwd UDI:** TCGTCGGCAGCGTCAGATGTGTATAAGAGACAG and **Rev UDI:** GTCTCGTGGGCTCGGAGATGTGTATAAGAGACAG). Each 20 µL reaction contained ∼2 µL of template DNA, 1×DreamTaq Green PCR Master Mix (Thermo Scientific), and 10 pmol of each primer. Cycling conditions were 94 °C for 3 min; 35 cycles of 94 °C for 30 s, 55 °C for 30 s, 72 °C for 60 s; and a final extension at 72 °C for 6 min. All samples were amplified in duplicate to ensure reproducibility. A second PCR was then performed to attach sample-specific barcodes (8-nt index sequences) using a universal adapter primer pair. Barcoded amplicons from each sample were pooled, purified, and normalized to ∼1-2 ng/µL. The library pool was sequenced on an Illumina MiSeq platform at Drexel Genomics Core Facility (2×250 bp paired-end reads). Raw reads were merged and filtered to remove primers/barcodes using FASTQ-join. High-quality reads were then clustered into haplotypes with SeekDeep software. For each isolate, we defined unique *Pv*msp1 haplotypes at a within-sample read frequency threshold ≥5%. Infections with multiple *Pv*msp1 haplotypes were classified as polyclonal, whereas those with a single haplotype were considered clonal (following the criteria of Lin *et al.*, 2015).

## Statistical analysis

Unless otherwise noted, statistical tests were two-sided. Paired comparisons used Wilcoxon signed-rank tests; group comparisons used ANOVA and/or Kruskal–Wallis tests; correlations used Pearson’s r; and haplotype enrichment used Fisher’s exact test. Multiple testing correction used Benjamini-Hochberg false discovery rate (FDR) threshold of 0.05 and log₂ fold change ≥1. Plots were generated in GraphPad Prism v11.0.0 (84), Qlucore Omics Explorer version 3.10, and Python version 3.x using standard scientific computing libraries including pandas, NumPy, scikit-learn, and matplotlib. Figures were assembled in Adobe Illustrator.

## Data availability

All sequence data generated in this study have been deposited in the NCBI Sequence Read Archive (SRA) under the BioProject PRJNA1328365 (see S3 table for SRA submission description). Code for preprocessing and analysis are available at https://github.com/Franck-Dumetz/Pv_Ethiopia.

## Acknowledgements

We sincerely acknowledge our dedicated field team in Jimma Town and Mizan Aman Hospital; Deje Lemese, Taye Teka, Meseret Teshome, Fikeret Legese, Wendemageng, Buzayehu, and Marta Zemede for their invaluable assistance in sample collection, processing, and organization. We are deeply grateful to the study participants, healthcare staff, and technicians from Jimma Town, Mizan Aman Hospital, and the surrounding communities for their cooperation and willingness to participate in this research. We also extend our appreciation to the Jimma University, Jimma Zone Health office and the collaborating health centers and hospitals for their continuous support; Regan Schroeder and Cheikh Dieng from Drexel University for their assistance in amplicon sequence data processing. We also thank Maryland and Drexel Genomics core for their support and expertise with Illumina sequencing.

## Author contributions

BRA, JP, DS, DY, and EL conceived and designed the study; BRA, TT, BL, and DT collected the samples; BRA, TT, and DT processed the samples; BRA, CF, FD, TS, DS, and EL analyzed the data; BRA, FD, JP, DY, DS, and EL wrote and edited the paper. All authors have read and approved the final manuscript.

## Supporting Information

### Supplementary Figure

**S1A-B Fig:**
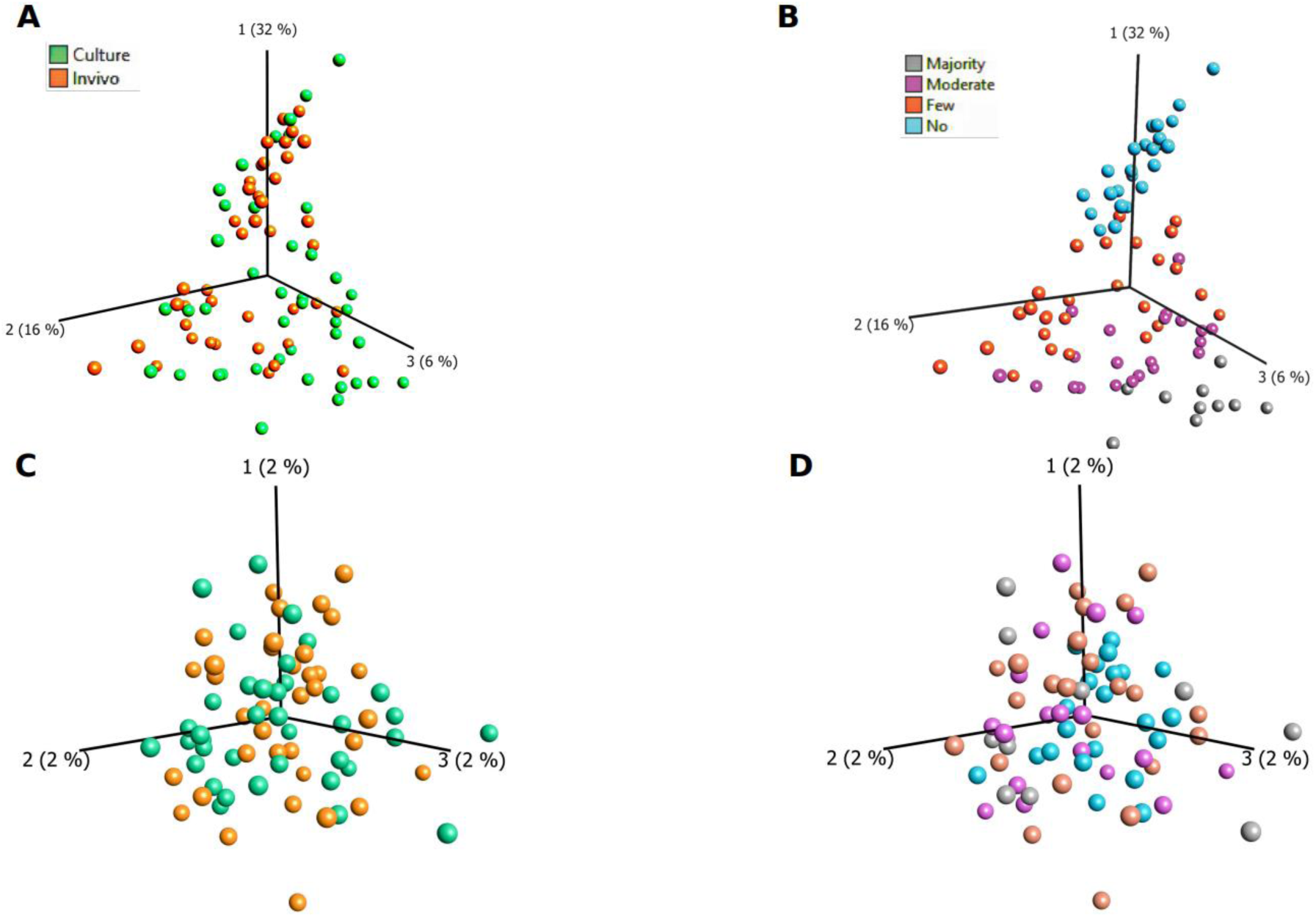
Unsupervised principal component analysis (PCA) revealed substantial overlap between *in vivo* and short-term cultured samples when colored by experimental condition. However, samples stratified by schizont stage fraction exhibited stage-associated clustering patterns, indicating that developmental composition is a primary driver of transcriptional variance. 1**C-D:** Following adjustment for parasite developmental stage, PCA showed no distinct clustering by Schizont stage composition.

### Supplementary Tables

S1 Table. Clinical and sequencing metadata for all analyzed samples including sample ID, infection source, culture condition, dominant haplotype, MOI classification, parasitemia category, and host Duffy genotype.

S2 Table. Parasite developmental stage fractions. Estimated stage proportions for each sample including ring, trophozoite, schizont, and gametocyte fractions.

S3 Table. Description of the SRA submission

### Supplementary Data

S1 Data. Raw gene expression matrix used for transcriptomic analyses.

S2 Data. Stage-adjusted residual expression values used for PCA, heatmap and Venn diagram.

S3 Data. Haplotype-level summary statistics for net haplotype change and enrichment analysis.

